# Operant alcohol self-administration targets GluA2-containing AMPA receptor expression and synaptic activity in the nucleus accumbens in a manner that drives the positive reinforcing properties of the drug

**DOI:** 10.1101/2024.09.13.612946

**Authors:** Sara Faccidomo, Briana L Saunders, Ashley M. May, Vallari R. Eastman, Michelle Kim, Seth M. Taylor, Jessica L. Hoffman, Zoé A McElligott, Clyde W Hodge

**Affiliations:** Bowles Center for Alcohol Studies School of Medicine, The University of North Carolina at Chapel Hill Chapel Hill, NC 27599 USA; Department of Psychiatry School of Medicine, The University of North Carolina at Chapel Hill Chapel Hill, NC 27599 USA; Department of Pharmacology School of Medicine, The University of North Carolina at Chapel Hill Chapel Hill, NC 27599 USA

**Keywords:** AMPA, GluA2, NSF, alcohol, self-administration, nucleus accumbens, basolateral amygdala

## Abstract

***Rationale*:** The positive reinforcing effects of alcohol (ethanol) drive its repetitive use and contribute to alcohol use disorder (AUD). Ethanol alters the expression of glutamate AMPA receptor (AMPAR) subunits in reward-related brain regions, but the extent to which this effect regulates ethanol’s reinforcing properties is unclear. ***Objective:*** This study investigates whether ethanol self-administration changes AMPAR subunit expression and synaptic activity in the nucleus accumbens core (AcbC) to regulate ethanol’s reinforcing effects in male C57BL/6J mice. ***Results:*** Sucrose-sweetened ethanol self-administration (0.81 g/kg/day) increased AMPAR GluA2 protein expression in the AcbC, without effect on GluA1, compared to sucrose-only controls. Infusion of myristoylated Pep2m in the AcbC, which blocks GluA2 binding to N-ethylmaleimide-sensitive fusion protein (NSF) and reduces GluA2-containing AMPAR activity, reduced ethanol-reinforced responding without affecting sucrose-only self-administration or motor activity. Antagonizing GluA2-lacking AMPARs, through AcbC infusion of NASPM, had no effect on ethanol self-administration. AcbC neurons receiving projections from the basolateral amygdala (BLA) showed increased sEPSC area under the curve (a measurement of charge transfer) and slower decay kinetics in ethanol self-administering mice as compared to sucrose. Optogenetic activation of these neurons revealed an ethanol-enhanced AMPA/NMDA ratio and significantly reduced paired-pulse ratio, suggesting elevated GluA2 contributions specifically within the BLA→AcbC pathway. ***Conclusions:*** Ethanol use upregulates GluA2 protein expression in the AcbC and AMPAR synaptic activity in AcbC neurons receiving BLA projections and enhances synaptic plasticity directly within the BLA→AcbC circuit. GluA2-containing AMPAR activity in the AcbC regulates the positive reinforcing effects of ethanol through an NSF-dependent mechanism, highlighting a potential therapeutic target in AUD.

## INTRODUCTION

The development of alcohol use disorder (AUD) represents maladaptive neuroplasticity, where alcohol hijacks neural mechanisms that regulate adaptive reward-seeking behavior, leading to excessive use (Hyman 2005; Kelley 2004a). Viewing AUD as aberrant plasticity in reward-associated neural circuits provides a compelling model for understanding how the memory of a drug’s rewarding properties is stored long-term, offering insights into the persistent nature of addiction-related behaviors (Nestler 2001). Understanding how drugs target specific plasticity mechanisms to produce and maintain harmful alcohol use can guide the development of novel medications (Anton et al. 1995; Gass and Olive 2008; Heilig and Egli 2006; Holmes et al. 2013; Hopf and Mangieri 2018; Kalivas and Volkow 2011).

Experience-dependent plasticity involves long-term increases in synaptic strength, largely due to upregulation of glutamate α-amino-3-hydroxy-5-methyl-4-isoxazolepropionic acid receptor (AMPAR) expression and activity at excitatory synapses (Malenka 2003; Malinow and Malenka 2002; Nicoll and Schulman 2023). AMPARs, formed by combinations of GluA1-4 subunits, are glutamate-gated ion channels. Their activity and role in plasticity are influenced by post-translational modifications such as phosphorylation, membrane trafficking, and subunit composition, with the presence or absence of GluA2 determining key functional properties related to memory induction, maintenance, and loss (Bredt and Nicoll 2003; Jiang et al. 2006; Malenka 2003).

Evidence suggests that experiences inducing plasticity, including exposure to ethanol and other drugs, promote an activity-dependent increase in synaptic expression of GluA2-lacking Ca^2+^-permeable AMPARs (CP-AMPARs) in brain regions regulating addiction-related behaviors, such as the nucleus accumbens (Acb) and amygdala (Bellone and Luscher 2006; Conrad et al. 2008; Faccidomo et al. 2021; Liu and Cull-Candy 2000; Mameli et al. 2011; Marty and Spigelman 2012; Wolf and Tseng 2012). Following plasticity induction, glutamate synapses stabilize and maintain memories via expression of GluA2-containing Ca^2+^-impermeable AMPARs (CI-AMPARs), which constitute about 95% of total AMPAR expression in the Acb, hippocampus (HPC), and prefrontal cortex (PFC) (Lu et al. 2009; Reimers et al. 2011). N-ethylmaleimide-sensitive factor (NSF) plays an essential role in the recruitment and insertion of these GluA2-containing AMPARS to the synaptic membrane and can regulate AMPAR GluA2 membrane trafficking (Song et al. 1998). Blocking synaptic trafficking of GluA2 with Pep2M, which disrupts the interaction between NSF and GluA2, inhibits long-term memory (Conboy and Sandi 2010; Migues et al. 2014).Thus, the dynamic expression and activity of specific AMPAR subunits may reflect the development, progression, and maintenance of addiction-linked maladaptive plasticity in brain function and behavior.

Growing evidence indicates that AMPAR activity in specific reward-related brain regions plays a crucial role in the positive reinforcing properties of ethanol (Acosta et al. 2011; Agoglia et al. 2015; Ary et al. 2012; Backstrom and Hyytia 2004; Bauer et al. 2022; Cannady et al. 2013; Cannady et al. 2016; Czachowski et al. 2012; Faccidomo et al. 2020; Hoffman et al. 2021; Hoffman et al. 2022; Hopf and Mangieri 2018; Montagud-Romero et al. 2021; Neasta et al. 2010; Pickering et al. 2007; Salling et al. 2016; Sciascia et al. 2015; Stephens and Brown 1999; Stuber et al. 2008). This activity establishes the neurobehavioral conditions for repetitive use and promotes the development and maintenance of AUD (Stolerman 1992). However, the role of specific AMPAR subunits in ethanol’s reinforcing effects is not fully understood. Our previous research investigated GluA2-lacking CP-AMPAR activity as a potential target of ethanol that regulates its positive reinforcing effects. Long-term ethanol self-administration by C57BL/6J mice, modeling the initial positive reinforcing effects in non-dependent subjects, increased phosphorylation of AMPAR pGluA1-S831, enhanced spontaneous excitatory postsynaptic currents (sEPSC), and promoted synaptic insertion of CP-AMPARs in basolateral amygdala (BLA) neurons projecting to the Acb (BLA→Acb) (Faccidomo et al. 2021). We also found that CP-AMPAR activity and synaptic trafficking of GluA1-containing AMPARs in the BLA are required for ethanol’s reinforcing effects (Faccidomo et al. 2021). These findings suggest that the initial development of ethanol’s reinforcing properties is mediated by plasticity at GluA1-containing AMPAR synapses in BLA neurons projecting to the Acb.

Despite these discoveries, the involvement of specific AMPAR subtypes in the Acb in ethanol’s reinforcing effects has not been evaluated. To address this gap in knowledge, this study evaluates the hypothesis that non-dependent ethanol self-administration targets AMPAR subunit expression and synaptic function in the Acb, regulating the drug’s reinforcing properties. The Acb, a central component of the brain’s reward circuitry, plays a pivotal role in mediating the reinforcing effects of drugs, including ethanol (Besheer et al. 2010; Hodge et al. 1995; Hodge et al. 1997; Hodge et al. 1992; Salling et al. 2017; Samson et al. 1999; Samson and Hodge 1993; Samson et al. 1992; Schroeder et al. 2008). Within the Acb, excitatory neurotransmission through ionotropic and metabotropic glutamate receptors is required for ethanol’s reinforcing and subjective stimulus properties (Besheer et al. 2003; Besheer et al. 2010; Besheer et al. 2009; Rassnick et al. 1992a; Rassnick et al. 1992b). The Acb receives glutamatergic projections from brain regions, including the BLA, where glutamate AMPA receptor activity is required for ethanol’s reinforcing effects (Agoglia et al. 2015; Cannady et al. 2016; Faccidomo et al. 2021; McCool et al. 2010; Salling et al. 2016), and evidence indicates that optogenetically enhanced excitatory transmission from the BLA to the Acb is rewarding (Stuber et al. 2011). Therefore, we also evaluated the impact of ethanol self-administration on AMPAR synaptic activity in Acb neurons receiving CaMKII-dependent excitatory projections from the BLA. Our findings indicate that the expression and synaptic activity of GluA2-containing CI-AMPARs in the Acb are influenced by ethanol and are necessary for its reinforcing properties.

## Materials and methods

### Mice

Adult (8-week-old) male C57BL/6J mice (total N=52) were purchased from Jackson Laboratories (Bar Harbor, Maine, US). Mice were group-housed (n=4/cage) in Plexiglas cages with food and water available ad libitum except during initial lever press training. As female mice are known to have different baseline levels of ethanol intake, and doubling the number of mice was outside the scope of the current study, additional experiments will need to be conducted to address potential sex differences that might affect ethanol regulation of GluA2-containing AMPA receptor expression and synaptic activity in the nucleus accumbens The mouse housing room and testing facility were maintained on a 12:12 light–dark cycle with lights off at 08:00 AM. Behavioral experiments were conducted during the dark to avoid light-associated disruption in mice (Roedel et al. 2006). Body weights were measured daily to calculate ethanol (EtOH) dosage consumed (g/kg). All experiments were approved by the Institutional Animal Care and Use Committee of the University of North Carolina at Chapel Hill and animals were cared for in accordance with the Guide for the Care and Use of Laboratory Animals (National Research Council (U.S.). Committee for the Update of the Guide for the Care and Use of Laboratory Animals. et al. 2011).

### Operant Ethanol Self-Administration Training and Baseline

Ethanol and sucrose-only self-administration studies were conducted in computer-controlled two-lever operant conditioning chambers (Med Associates, Fairfax, VT USA) as previously described (Faccidomo et al. 2009; Faccidomo et al. 2020; Faccidomo et al. 2016; Faccidomo et al. 2015; Salling et al. 2008; Salling et al. 2016; Salling et al. 2017). Briefly, lever-press responding was trained and maintained by sucrose-only (2% w/v; parallel non-drug control) or sweetened ethanol (9% v/v ethanol + 2% w/v sucrose) on a fixed-ratio 4 (FR4) schedule of reinforcement. Each chamber was equipped with a syringe-pump liquid delivery system and two response levers. Responses on an “active” lever (randomized across subjects) triggered the pump to deliver 0.014 ml of the reinforcing solution into a liquid cup receptacle equipped with a photobeam head-entry detector to allow counting of headpokes. At the end of each session, an experimenter checked the cups to verify consumption. Ethanol intake (g/kg) was calculated from the volume delivered / consumed. Responses on an “inactive” lever were recorded but produced no programmed consequences. After daily 1-hr baseline sessions (40-45 sessions), separate groups of mice were used for 1) measurement of AMPAR subunit expression via immunoblot; 2) whole-cell patch-clamp recordings of BLA→ nucleus accumbens core (AcbC) projection neurons; or 3) site-specific infusion of AMPAR inhibitor compounds targeting the AcbC. Mice that did not exhibit baseline levels of operant behavior ≥10 reinforcements/hr were excluded.

### Gel Electrophoresis and Immunoblot

Immediately after the 40^th^ self-administration session, mice (n=8/group) were rapidly decapitated, and brains were flash frozen in −20°C isopentane and stored at −80°C until use. Bilateral tissue punches (1 x 1 mm circle; **Fig 1C**) taken from Acb, primarily AcbC possible inclusion of minimal dorsal Acb shell, collected from frozen tissue on a cryostat (Leica Biosystems, Deer Park, IL USA) and homogenized in an SDS homogenization buffer containing protease/phosphatase inhibitors. Protein concentration was quantified using a Bradford colorimetric assay kit (BCA Assay; Pierce Protein Biology, ThermoFisher Sci) and gel electrophoresis and immunoblots were conducted as previously reported (Agoglia et al. 2015; Faccidomo et al. 2018; Wilkie et al. 2007). Briefly, sample protein (5-8 µg) and molecular weight ladders were loaded onto a 4–15% Criterion TGX precast gels (Bio-Rad) for gel electrophoresis separation. Proteins were transferred to PVDF membranes using an iBlot® Semi-Dry blotting system (Thermo Fisher Sci).

**Figure 1.**
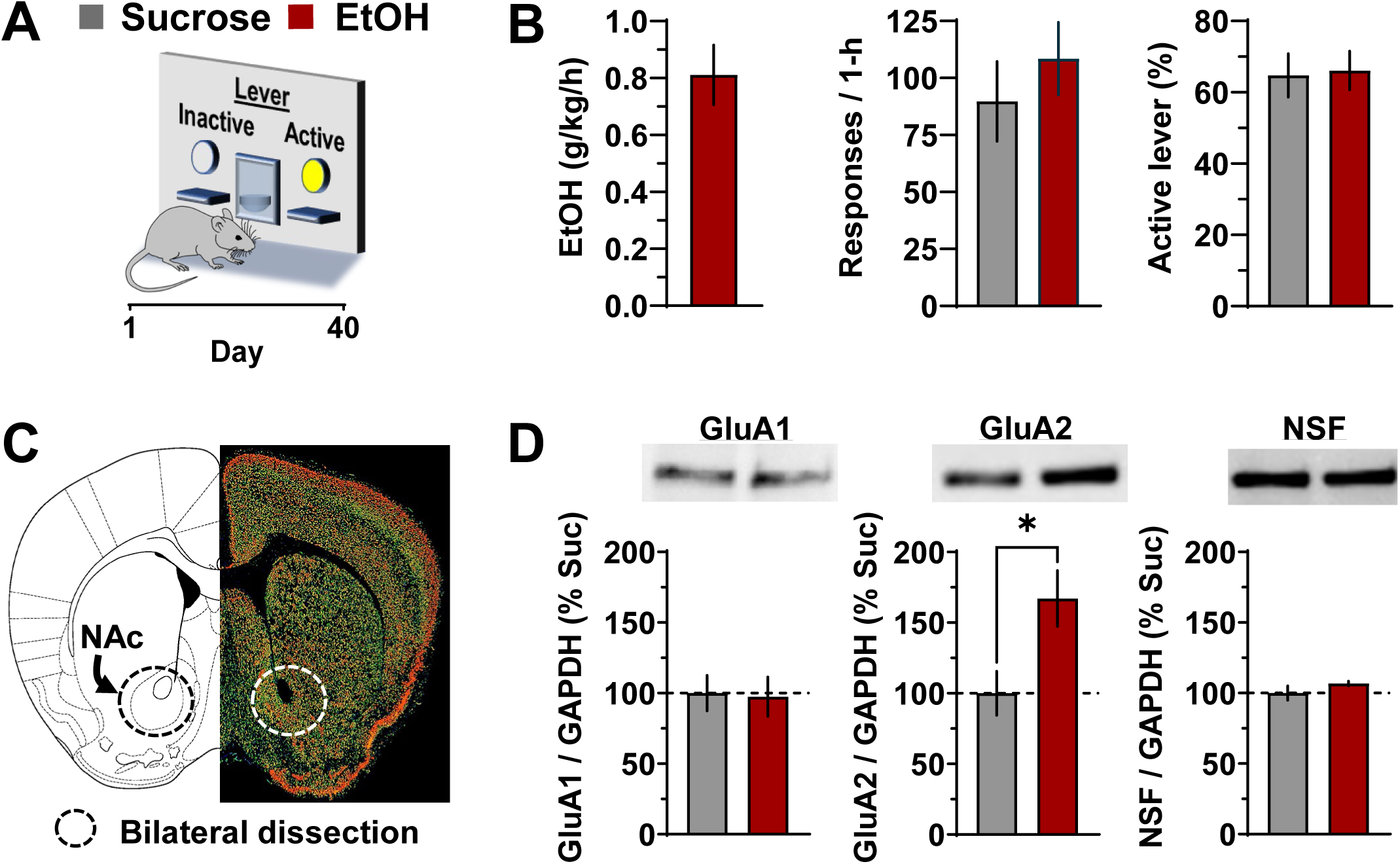
Operant ethanol self-administration increases AMPAR GluA2 subunit protein expression in the AcbC. **(A)** Schematic of operant conditioning chamber showing active and inactive lever with timeline. (**B**) Measures of operant ethanol (EtOH) and sucrose self-administration plotted as MEAN±SEM for n=8 mice per group. (**C**) Mouse brain atlas image juxtaposed with photomicrograph showing GluA2 gene expression in mouse brain (Allen Brain Atlas) with the dissection region of the AcbC highlighted by the dashed-line circle. (**D**) Ethanol self-administration (red bars) resulted in a significant increase in GluA2 protein expression in the AcbC but no change in GluA1 or NSF; * -indicates P<0.05 relative to sucrose (gray bars) control. Immunoblot data represent MEAN±SEM from n=5-8 mice per condition.

For protein measurement with immunoblots, a two-day protocol was used. Membranes were rocked in a blocking buffer (5% Normal Goat Serum [NGS]; Vector Labs) for 2 hrs at room temperature and then incubated with the following primary antibodies overnight at 4°C on separate blots: monoclonal mouse anti-GluA2 (1:2000, NeuroMab; #75-002), monoclonal rabbit anti-GluA1 (1:1000, Millipore; #04-855), polyclonal rabbit anti NSF (N-ethylmaleimide-sensitive fusion protein (NSF); 1:5000, Cell Signaling; #3924). The next day, membranes were washed and incubated with an HRP-conjugated secondary antibody (goat anti-rabbit and goat anti-mouse, 1:10,000; Jackson Immuno Research) for 1 hr. The chemiluminescent signal was visualized with ECL Select or Prime (GE Healthcare) and digitally captured using a digital imager (ImageQuant LAS 4000, GE Healthcare). On a different day, membranes were probed with an anti-mouse GAPDH antibody (1:10,000, Advanced Immunochemical) which served as a non-specific loading control. The optical density of each band was visualized and quantified using ImageQuant TL software. The optical density of target bands was normalized to the optical density of GAPDH and expressed as a ratio (target/GAPDH). To standardize data presentation across multiple gels, in addition to normalizing the optical density of target bands to GAPDH, data ratios within each gel were expressed as a percentage relative to the mean value from the sucrose group (parallel non-drug control group). This additional step also allows for accurate comparison of relative protein expression changes between the sucrose and ethanol groups. Each blot contained all samples from both the sucrose and ethanol groups and significant differences in pre-normalization raw values not found.

### Stereotaxic Surgery

Operant trained mice underwent stereotaxic surgery for bilateral guide cannula implantation into the AcbC (AP: +1.6, ML: +/-.75, DV: −3.8 mm from skull; 26 gauge, 6mm length, 1.5 C-C distance; Plastics One/Protech International) or bilateral AAV injections with AAV2-CaMKIIa-hChR2(H134R)-EYFP (Deisseroth Stock Vectors from UNC Gene Therapy Vector Core) into the BLA (AP: −1.3, ML: +/-3.4, DV: −4.4 mm from dura). Microinjection cannula surgeries occurred after 40 sessions of acquisition so the cannulae stayed patent throughout the microinjection study. Surgeries for the AAV infusion occurred before acquisition of operant responding to give time to the vector to express and travel from BLA to AcbC such that electrophysical recordings could be occur after 40-45 days of operant ethanol or sucrose self-administration sessions. Briefly, mice were anesthetized with 2-4% isoflurane throughout the duration of the aseptic surgery. The skull was visualized, Bregma was determined and (1) guide cannula was stereotaxically placed targeting the AcbC and affixed with dental cement or (2) the ChR2 AAV was bilaterally infused to the BLA at a rate of 0.1 ul/min (0.3 ul/side). Next, the surgical opening was closed, and mice were given a single bolus saline (1ml, s.c.) and ibuprofen (15mg/kg, 3days) to aid in recovery from surgery.

### Intra-AcbC Microinjections

*Operant self-administration*. Mice were returned to daily operant sessions 1 week after recovery from surgery. Baseline responding was re-established, and sham control injections were conducted to habituate the mice to the microinjection procedure. For drug microinjections, injectors (bilateral, 7mm length, 1mm past cannula guide) were inserted into the guide cannulae and NASPM (0, 1, or 10 µg/0.5ul/side) or Pep2m (0, 1, or 10 µg/0.5ul/side) was infused at a flow rate of 0.125 µl/min for 4-min in separate groups of mice. Injectors were left in place for an additional 1-min to allow drug diffusion. Mice were then placed immediately in the operant chambers for 1-hr self-administration sessions. Drug doses were administered according to a within-subject Latin square design to control for potential dose-order effects. Each injection was administered only after operant responses returned to baseline, with no more than two injections per week and at least 48 hours between each injection. No carryover effects of either drug were detected on days following the injections. Bilateral injection sites in the AcbC were confirmed histologically.

*Locomotor activity and thigmotaxis*. To address the possibility that changes in ethanol self-administration induced by intra-AcbC NASPM or Pep2m infusion were associated with nonspecific behavioral effects, spontaneous open-field locomotor activity (general motor function) and thigmotaxis (anxiety-like behavior) were assessed in 1-hr tests in the same mice after completion of the self-administration dose-effect curves as reported previously (Agoglia et al. 2016; Faccidomo et al. 2015; Hodge et al. 2004; Hodge et al. 1999; Hodge et al. 2006; Hodge et al. 2002).

### Whole-cell patch-clamp recordings

Slices (300 µm) containing the Acb were prepared as previously described (Faccidomo et al. 2021). Briefly, mice were anesthetized with isoflurane, and brains were extracted and sliced on a Leica VT1200 or VT1000 in ice cold high-sucrose low Na^+^ artificial cerebral spinal fluid (sucrose ACSF, in mm: 194 sucrose, 20 NaCl, 4.4 KCl, 2 CaCl_2_, 1 MgCl_2_, 1.2 NaH_2_PO_4_, 10 glucose, 26 NaHCO_3_) that had been perfused for at least 15 minutes in carbogen (95% O_2_, 5% CO_2_). Subsequent to slicing, sections containing the nucleus accumbens equilibrated in the same ACSF that was used for recordings (in mm: 124 NaCl, 4.4 KCl, 2 CaCl_2_, 1.2 MgSO_4_, 1 NaH_2_PO_4_, 10 glucose, 26 NaHCO_3_, 95% O_2_, 5% CO_2_34°C) with picrotoxin (25 µM) for 30 minutes prior to being transferred to the recording chamber where they were perfused (2 ml/min) with ACSF for an additional 30 minutes prior to recording. All recordings were made in Cs-gluconate intracellular solution (in mM: 135 Cs-gluconate, NaCl 5, MgCl_2_ 2, HEPES 10, EGTA 0.6, Na_2_ATP 4, Na_2_GTP 0.4). All recordings were made in ClampEx 10 and were analyzed with ClampFit 10 (Molecular Devices), or MiniAnalysis (Synaptosoft). Spontaneous EPSCS were recorded at −80 mV, paired-pulse ratios were recorded at −70 mV, and AMPA/NMDA ratios were recorded at −70/+40 mV. Optogenetic stimulation of ChR2 was performed with a 470 nm LED (CoolLabs). Evoked excitatory post-synaptic currents (eEPSCs) were evoked by local stimulation with Ni-chrome bipolar electrodes while neurons were voltage-clamped at −70 mV and +40 mV to determine AMPA/NMDA ratios. Amplitudes for both optically and electrically evoked NMDA currents were recorded 50 ms following the stimulation. Drugs were bath applied at final concentrations.

### Drugs

NASPM [N-[3-[[4-[(3-Aminopropyl)amino]butyl]amino]propyl]-1-naphthaleneacetamide trihydrochloride; Bio-Techne, Tocris, USA) a selective GluA2-lacking CP-AMPAR antagonist, was dissolved in phosphate-buffered saline for bath application in patch-clamp electrophysiological experiments or artificial cerebrospinal fluid (ACSF) for site-specific infusion in the AcbC in behavioral procedures. Myristoylated Pep2m (Pep2m; Formula: C_63_H_118_N_18_O_14_S; Sequence: KRMKVAKNAQ; Bio-Techne, Tocris, USA) a cell-permeable inhibitor of GluA2-containing AMPARs that acts via inhibition of the interaction between the GluA2 C-terminus and N-ethylmaleimide-sensitive fusion protein, was dissolved in aCSF for site-specific inhibition of GluA2-containing AMPARs in the AcbC of behaving mice. Drug doses were selected to be in the range used in prior studies that evaluated site-specific effects of Pep2M (Conboy and Sandi 2010; Migues et al. 2014) or NASPM on behavior (Faccidomo et al. 2021; Martinez-Rivera et al. 2017) and synaptic activity (Coombs et al. 2023; Faccidomo et al. 2021; McElligott et al. 2010).

### Statistical Analysis

Data are presented as MEAN±SEM. All statistical analyses and graphical representations of data were performed using GraphPad Prism software (version 10.3.1, GraphPad Software, San Diego, CA). Group comparisons were conducted via unpaired t-test, one-way RM ANOVA, or two-way RM ANOVA. Post hoc comparisons were conducted with Dunnett’s or Šídák’s multiple comparison procedures, where appropriate. P values of < 0.05 were statistically significant.

## Results

### Self-administered ethanol increases GluA2 protein expression in the Acb

We first sought to determine if non-dependent operant ethanol self-administration is associated with altered AMPAR subunit expression in the AcbC. Parallel groups of male C57BL/6J mice were trained to self-administer sweetened ethanol (n=8) or sucrose-only (n=8) during 40 daily 1-hr sessions (**Fig 1A & B**). Importantly, analysis of operant self-administration behavior showed no statistically significant group differences on measures of response rate (responses / 1-hr) or the percentage of responses on the active lever (**Fig 1B**) indicating that differences observed in protein expression are not attributable to differential reinforcer efficacy, motor activity, learning and memory, or general reward processing between sucrose and ethanol trained mice. The dosage of ethanol consumed was *X̅* = 0.81 g/kg/1hr.

Western blot analysis was conducted on whole-cell lysates that targeted the AcbC where *Gria2* gene expression is higher than the adjacent ventral shell subregion (**Fig 1C; Allen Brain Atlas**). Immunoblot analysis of protein expression found that ethanol self-administration produced no change in AMPAR GluA1 subunit expression (**Fig 1D, left panel**). However, there was a statistically significant increase in AMPAR GluA2 subunit protein expression in ethanol self-administering mice as compared to sucrose administering controls (**Fig 1C, middle panel;** t_9_ = 2.6, P = 0.029). We also showed that ethanol had no immediate effect on NSF expression in the AcbC as compared to sucrose (**Fig 1C, right panel**) indicating that the potential for NSF-dependent anchoring of GluA2-containing CI-AMPARs to AcbC excitatory synapses remained intact. This suggests that a history of non-dependent ethanol self-administration is only associated with an increase in GluA2-containing AMPARs in the AcbC.

### GluA2-containing AMPAR activity in the AcbC is required for the positive reinforcing effects of ethanol

This experiment sought to determine if GluA2-containing AMPARs in the AcbC mechanistically regulate the positive reinforcing effects of ethanol. To address this question, C57BL/6J mice received site-specific injections of myristoylated Pep2m targeting the AcbC prior to operant ethanol (n=8) or sucrose (n=9) self-administration sessions (**Fig 2A)**. Two-way RM-ANOVA identified a significant main of time (F_12,84_ = 158, P<0.0001) on the rate of ethanol reinforced responding. There was also a significant time x Pep2m interaction (F_24,168_ = 2.708, P=0.0001) but no main effect of Pep2M alone (F_2,14_ = 2.99, P=0.=.082). Accordingly, Dunnett’s multiple comparison analysis of the interaction showed that effects of Pep2m (10 µg) emerged over time with significant decreases in EtOH-reinforced response rate occurring from 20 – 60 minutes of the 1-hr session (**Fig 2B**). This indicates that self-administration behavior began at a normal rate but was reduced after 15-min of EtOH access, suggesting no general disruptions in the onset of operant behavior. One-way RM-ANOVA of total session results showed that the Pep2M-induced reduction in EtOH-reinforced response rate was associated with an overall significant dose-dependent decrease in the number of reinforcers earned (F_2,20_ = 3.9, P = 0.037). Dunnett’s multiple comparisons test showed that effects of Pep2M were specific to the 10-µg dosage (**Fig 2C**). Pep2m had no effect on general consummatory behavior as inferred by no change in the number of headpokes in the liquid delivery cup per reinforcer (F_2,13_ = 0.33, P = 0.723; **Fig 2D**).

**Figure 2.**
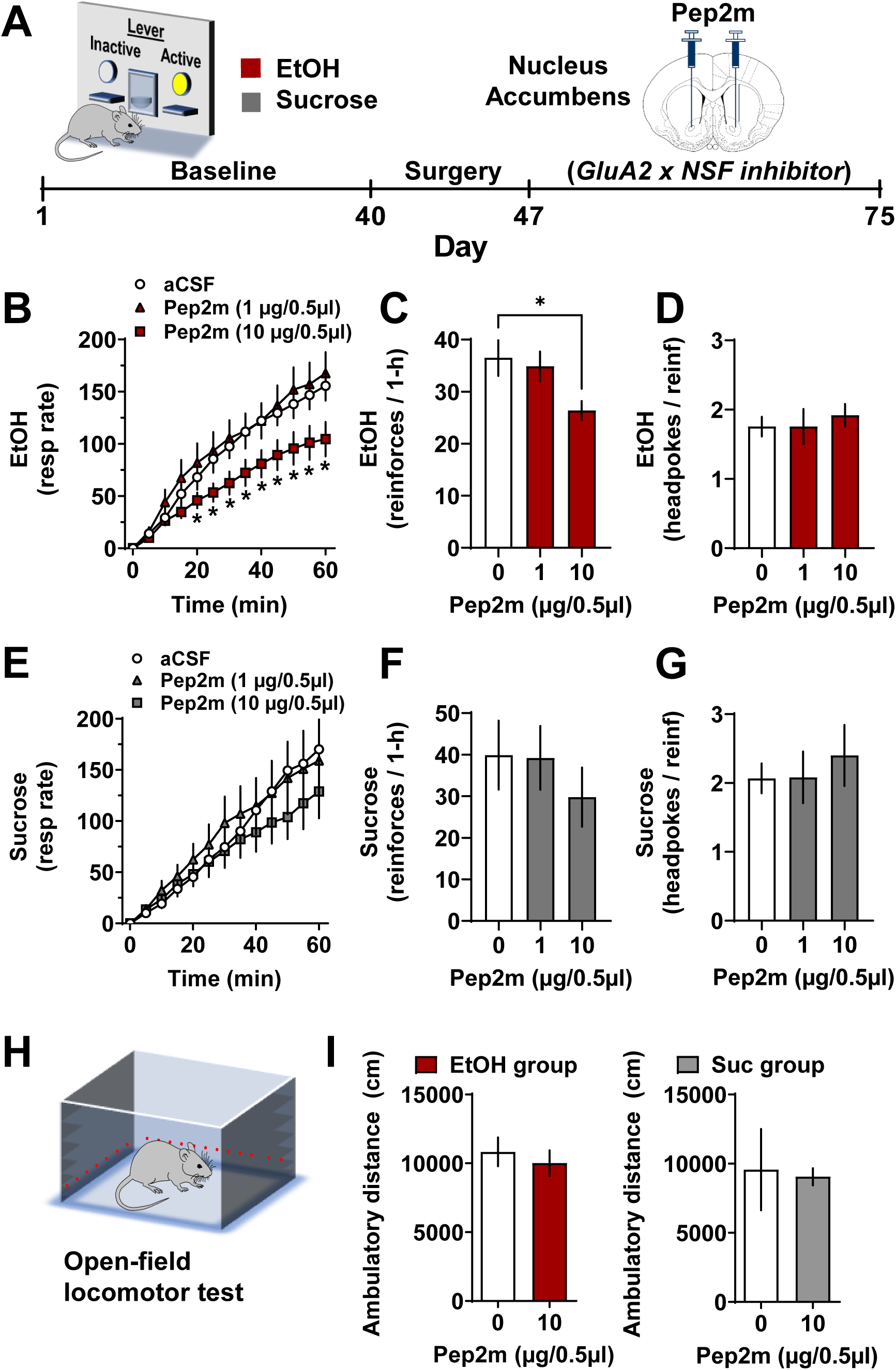
GluA2-containing AMPAR activity in the AcbC is required for the positive reinforcing effects of ethanol. (**A**) Schematic showing timeline (days) of baseline behavioral training with ethanol (EtOH, n=8) and sucrose-only (n=9) reinforcement, surgery, and Pep2m infusion targeting the AcbC. (**B**) Line graph shows EtOH-reinforced response rate (resp / 5-min) plotted as a function of time (min). * -indicates that Pep2m (10 µg) significantly decreased EtOH-reinforced response rate as compared to vehicle control (Pep2m 0 µg) at corresponding time points; Dunnett’s multiple comparison test, P<0.05. (**C**) Number of ethanol reinforcers earned during total session time (1-hr) plotted as a function of Pep2m dosage. * -indicates significant difference from vehicle control; Dunnett’s multiple comparison test, P<0.05. (**D**) Head pokes per ethanol reinforcer delivery were unchanged by Pep2m infusion. (**E-G**) Parameters of sucrose-only self-administration were not changed by infusion of Pep2m in the AcbC. (**H**) Schematic of open-field locomotor activity monitor (**I**) Pep2m (0 or 10 µg) infusion in the AcbC had no effect on locomotor activity in the ethanol or sucrose-only groups. All data are MEAN±SEM.

Self-administration of sucrose in a parallel behavior-matched control group (n=9) showed a predicted main effect of time on sucrose-reinforced response rate (F_1.1,8.8_ = 22.44, P = 0.001). However, there was no significant effect of Pep2m (F_1.3,10.8_ = 1.28, P = 0.30) and no interaction (F_2.3,18.6_ = 2.15, P = 0.14; **Fig 2E**). Analysis of total session data showed that Pep2m had no effect on the number of reinforcers earned per 1-hr (F_2,16_ = 2.3, P = 0.13; **Fig 2F**) or on the number of headpokes per reinforcer (F_2,16_ = 0.79, P = 0.47; **Fig 2G**). Moreover, infusion of the effective dosage of Pep2m (10 µg/0.5µl), that reduced operant ethanol self-administration, had no effect on locomotor activity by either ethanol (t_8_ = 0.7, P = .25) or sucrose (t_8_ = 1.56, P = 0.08) mice in separates tests conducted after completion of self-administration dose-response curves (**Fig 2H & I**), indicating the absence of nonspecific effect on motor ability. These results show that GluA2-containing AMPAR activity in the AcbC is required for the full expression of the positive reinforcing effects of ethanol.

To test the specificity of GluA2-containing AMPARs in the functional regulation of ethanol reinforcement, we injected NASPM (0, 1 or 10 µg/0.5µl) into the AcbC of a separate group of ethanol self-administering mice (n=6) to inhibit GluA1-containing CP-AMPARs (**Fig 3A**). EtOH-reinforced response rate increased as a function of time during 1-hr sessions (F_12,60_ = 126, P < 0.0001); however, there was no effect of NASPM (F_2,13_ = 0.33, P = 0.723) and no time x NASP interaction (F_24,107_ = 0.33, P = 0.723; **Fig 3B, left**). Accordingly, there was no effect of NASPM on the number of EtOH reinforcers earned (F_2,10_ = 0.13, P = 0.982; **Fig 3B, middle**) or number of headpokes per reinforcer (F_2,10_ = 2.24, P = 0.16; **Fig 3B, right**). In addition, there was no significant effect on locomotor activity in the open-field test following NASPM (0 or 10 µg/0.5µl) infusion in the AcbC (t_14_ = 1.6, P=0.07; **Fig 3C**). This finding suggests that CP-AMPARs in the AcbC do not regulate the reinforcing effects of ethanol and suggest specificity of control by GluA2-containing AMPARs.

**Figure 3.**
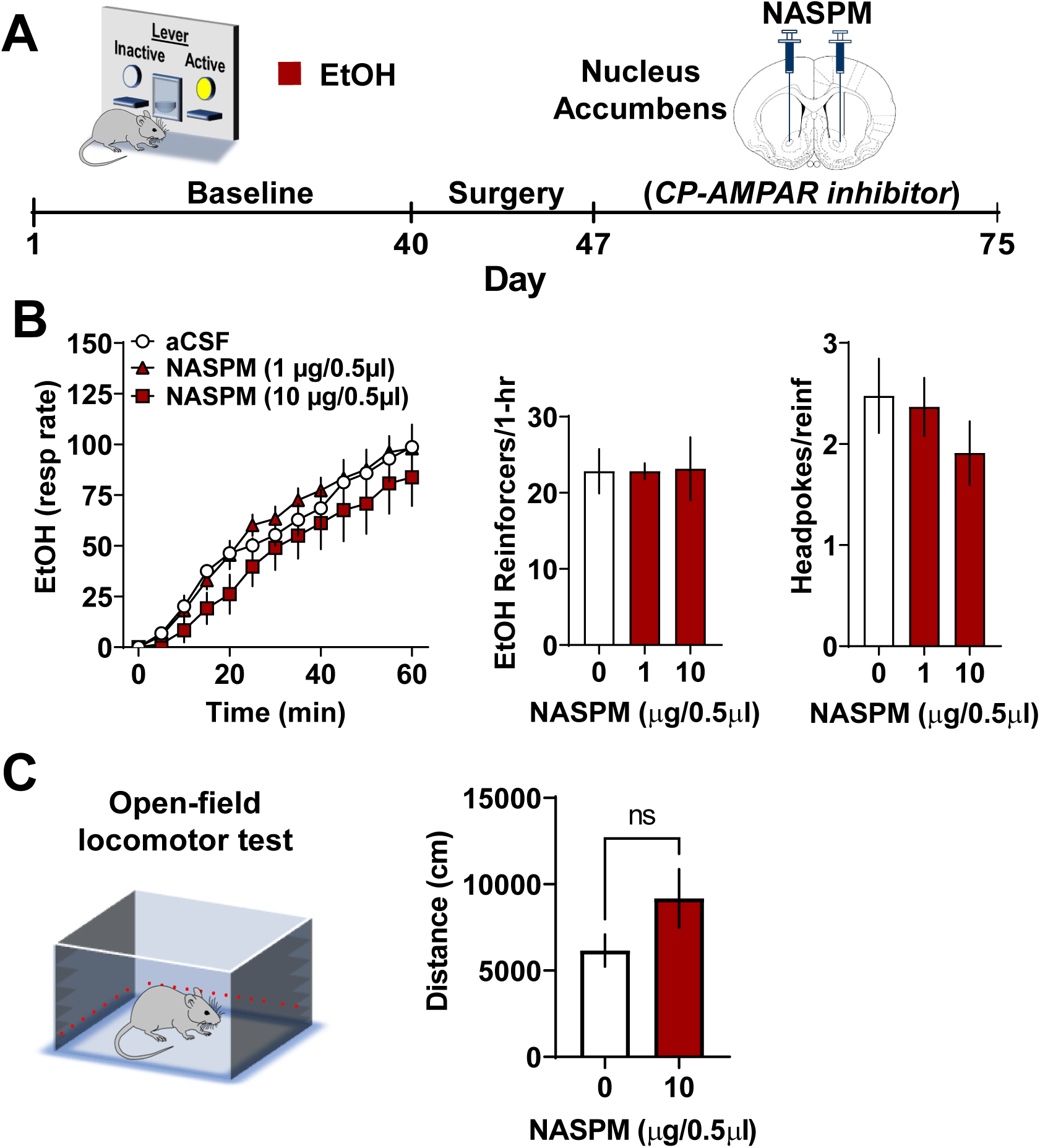
GluA2-lacking CP-AMPAR activity in the AcbC does not regulate the positive reinforcing effects of ethanol. (**A**) Schematic showing timeline (days) of baseline ethanol (EtOH) self-administration training (n=7), surgery, and NASPM infusion targeting the AcbC. (**B**) NASPM had no effect on ethanol reinforced response rate, number of reinforcers earned, or headpokes in the delivery cup. (**C**) Schematic of open-field locomotor test chamber. (**D**) Ambulatory distance traveled in the open field after NASPM administration showing no effect. All data are MEAN±SEM.

### Self-administered ethanol increases synaptic activity of CI-AMPARs in AcbC neurons receiving BLA projections

We discovered previously that operant ethanol self-administration selectively increased synaptic activity and insertion of GluA2-lacking CP-AMPARs in the BLA of neurons that project to the Acb, as compared to behavior-matched sucrose controls (Faccidomo et al. 2021). However, that study did not evaluate the BLA to Acb circuit. Thus, to determine if the ethanol-induced upregulation of AMPAR GluA2 protein expression is associated with altered synaptic activity in AcbC neurons receiving BLA projections, male C57BL/6J mice received site-specific infusion of AAV2-CaMKIIa-hChR2(H134R)-EYFP in the BLA to probe neurons in the AcbC receiving BLA synaptic input. After a 7-day recovery, mice were separated into two groups and trained to self-administer sweetened ethanol (n=7) or sucrose-only (n=6). After a 43d baseline and 1-d of ethanol clearance, brains were removed for patch-clamp electrophysiological recordings of electrical-or light-evoked ChR2-positive neurons in the AcbC (**Fig 4A)**. Infusion of ChR2 in the BLA under the CaMKII promoter resulted in significant expression in AcbC / shell subregions (**Fig 4B**). Behavioral results showed that mice self-administered ethanol and sucrose (control) solutions at an equivalent rate indicating no potential impact of differential behavior between the groups (Responses / 1-hr: t_11_ = 1.17, P = 0.27; Active lever %: t_11_ = 0.81, P = 0.44; Headpokes / 1-hr: t_11_ = 1.31, P = 0.22) (**Fig 4C**). The MEAN±SEM dosage of EtOH self-administered was 0.79±0.09 g/kg/1-hr (**Fig 4C**).

**Figure 4.**
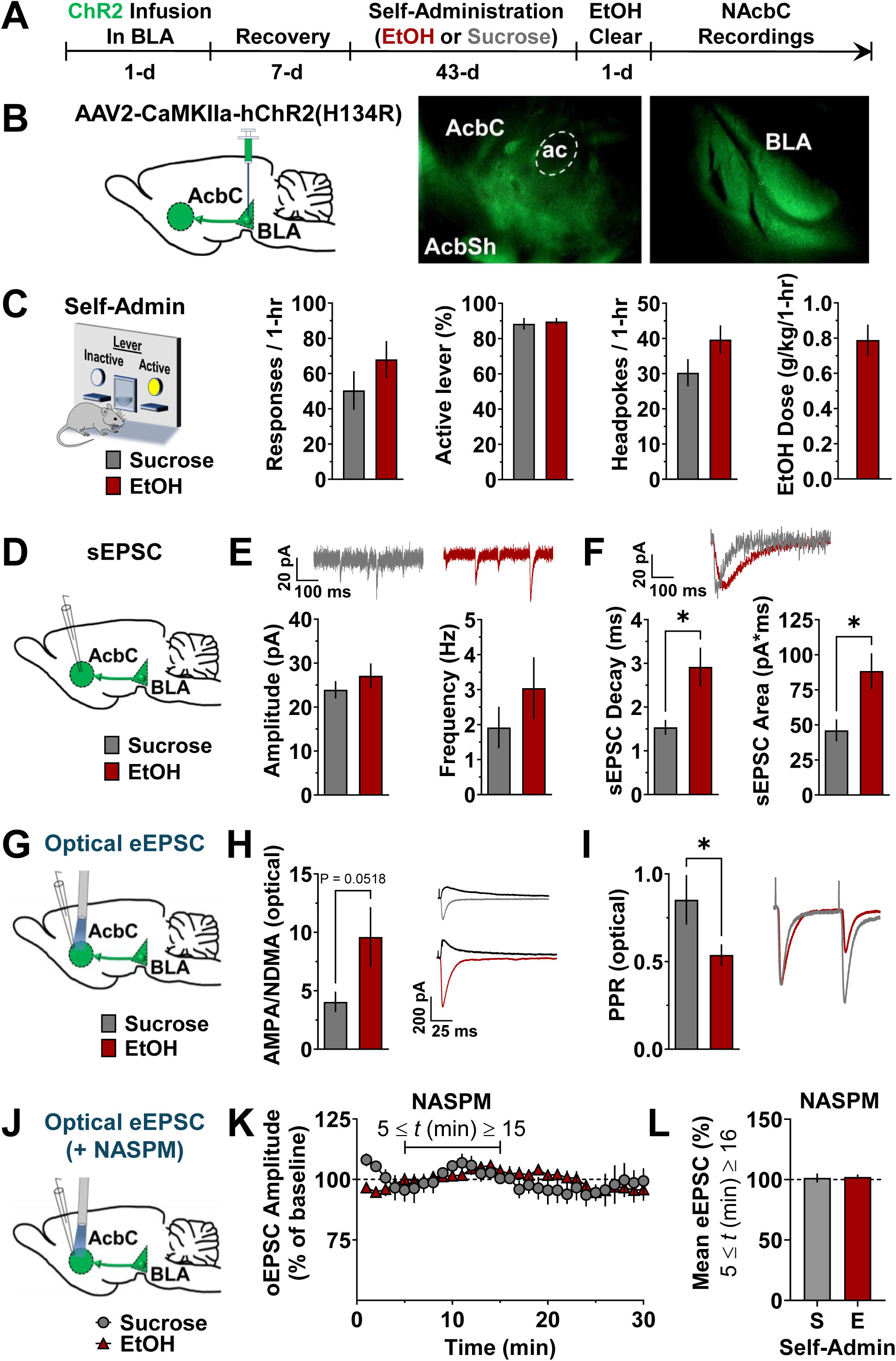
Ethanol self-administration increases CI-AMPAR but not CP-AMPAR activity in AcbC neurons receiving input from the BLA. (**A**) Experimental timeline (days) and schematic showing ChR2 infusion in BLA followed by recovery and 43-days of ethanol (EtOH, n=8) or sucrose (Suc, n=6) self-administration. BLA recordings were conducted 24-h after the last self-administration session. (**B**) Schematic of ChR2 infusion in mouse BLA and photomicrograph showing expression in the BLA and AcbC. (**C**) Parameters of ethanol (red bars) and sucrose (gray bars) self-administration show behavior-matched performance (n=6-8 per group). (**D**) Schematic illustrating sEPSC in AcbC neurons from BLA projections. (**E**) Average plots showing no change in sEPSC amplitude or frequency in AcbC, (**F**) decay time (ms), and area under the curve in EtOH self-administering mice as compared to sucrose-only controls. * -indicates significantly different from sucrose; t-test, P<0.05. Representative traces illustrating EtOH-induced increases are shown above and to the right of average plots. (**G**) Schematic illustrating optically evoked EPSC (oEPSC) in AcbC neurons receiving BLA projections. (**H**) Plots showing a strong trend for an increase in optically evoked AMPA/NMDA ratio (p = 0.052) and (**I**) a significant reduction in the paired-pulse ratio (PPR). * -indicates significantly different from sucrose-only controls; t-test, P<0.05. (**J**) Schematic showing oEPSC in AcbC neurons receiving BLA projections following bath application of NASPM. (**K**) Time course of optical eEPSC amplitude (MEAN±SEM) during baseline (min 1 – 4) and bath application of NASPM (100 µM; min 5 - 16) from sucrose vs. ethanol self-administering mice. Data are normalized to minute 4 of the baseline. (L) Bar graph shows summary of eEPSC during NASPM bath for mice with a history of sucrose (S) and ethanol (E) self-administration.

Electrophysiological recordings from AcbC neurons revealed that EtOH self-administration had no effect on sEPSC amplitude (t_18_=0.89, P = 0.3878) or sEPSC frequency (t_18_=1.014, P = 0.326) as compared to sucrose-only controls (**Fig 4D-E**). Interestingly, however, we found that ethanol self-administration significantly slowed sEPSC decay kinetics as measured by decay time (t_18_=2.7, P = 0.012) and significantly increased the area under the curve (t_18_=2.6, P = 0.015; **Fig 4F**), which is consistent with an increase in glutamatergic transmission and upregulated GluA2-containing AMPARs in the AcbC (Chater and Goda 2014; Pampaloni and Plested 2022).

Optically evoked EPSC (oEPSC) in AcbC neurons (**Fig 4G**) showed a strong trend for an increase in AMPA/NMDA ratio (t_22_=2.1, P = 0.0518) (**Fig 4H**). In addition, there was a significant reduction in the paired-pulse ratio t_22_=2.4, P = 0.02), together these results suggest that a history of ethanol self-administration is associated with upregulated AMPAR synaptic activity, and that ethanol use is associated with an enhancement of glutamate release in AcbC neurons receiving BLA projections (**Fig 4I**). Surprisingly, there was no ethanol-associated change in the AMPA/NMDA ratio when measured after local electrical stimulation of nonspecific afferents in the AcbC (MEAN±SEM: sucrose 6.09±1.6; ethanol 8.6±3.42, t_15_=0.57, P=0.58), suggesting specificity of the BLA projection to the AcbC.

Since the difference between the optically evoked AMPA/NMDA ratio for sucrose and ethanol showed a strong trend for statistical significance, we calculated effect size metrics (Hedges’ *g* and effect-size *r*) and 95% confidence intervals (CIs) to provide more information regarding the magnitude and strength of the relationship. Hedges’ g = (9.602 −4.046) ⁄ 9.382455 = 0.592169 suggests a moderate effect size, indicating that the ethanol condition may elevate the AMPA/NMDA ratio compared to sucrose. This is further supported by an effect-size *r* of 0.325, indicating that a meaningful portion of the variability in the AMPA/NMDA ratio is due to differences between sucrose and ethanol groups. Examination of the 95% CIs supports this interpretation, with the sucrose mean AMPA/NMDA ratio ranging from 2.030 to 6.062 and the ethanol mean ratio ranging from 4.234 to 14.97. The lack of overlap between the lower bound of the ethanol CI and the upper bound of the sucrose CI suggests that ethanol influences the AMPA/NMDA ratio, despite the marginal t-test p-value.

Although the present results are consistent with upregulated GluA2-containing CI-AMPAR activity (protein expression, decay kinetics and PPR), our previous findings indicate that ethanol self-administration increased synaptic insertion of CP-AMPARs in the BLA (Faccidomo et al. 2021). Thus, we assessed the potential contribution of GluA2-lacking CP-AMPARs to synaptic transmission in AcbC neurons receiving BLA projections by bath application of the CP-AMPAR blocker NASPM and evaluated oEPSC (**Fig 4J**). Two-way RM-ANOVA (EtOH x time) conducted on the total timecourse of exposure found that NASPM (100 µM) had no effect on oEPSC amplitude expressed as a percentage of baseline in ChR2-positive neurons (EtOH: F_1,9_ = 0.52, P = 0.49; Time: F_29,261_ = 1.25, p = 0.18; EtOH x Time: F_29,261_ = 0.93, P=0.56; **Fig 4K**). NASPM also had no effect on the amplitude of oEPSCs when analyzing percent change from the interval before bath application (t22 = 0.66, P = 0.52; **Fig 4L**). Lack of NASPM effects, combined with the change in the decay of the sEPSCs and decrease in PPR, suggests that the EtOH-induced changes in excitatory transmission are not a function of synaptic insertion of CP-AMPARs but may reflect ethanol-induced upregulated GluA2-containing CI-AMPAR activity.

## Discussion

Through a series of experiments combining operant ethanol self-administration with molecular, physiological, optogenetic, and pharmacological approaches, we found that GluA2-containing CI-AMPARs in the AcbC are targeted by self-administered ethanol. This mechanism underlies the drug’s positive reinforcing properties, which are essential for the experience-dependent development and progression of AUD (Wise and Koob 2013). This discovery deepens our understanding of alcohol addiction’s molecular mechanisms and neural circuitry. Given GluA2’s role in memory maintenance, it also unveils a potential therapeutic target for medications aimed at promoting the extinction of behavioral pathologies associated with AUD, including chronic alcohol drinking.

### Self-administered ethanol increases GluA2 protein expression in the AcbC

A novel finding of this study is that chronic operant self-administration of sweetened ethanol significantly increased AMPAR GluA2 subunit protein expression by 67% in the AcbC relative behavior-matched sucrose controls. Importantly, no group differences were seen in AMPAR GluA1 protein expression in the AcbC. We measured NSF expression due to its crucial role in recruiting and inserting GluA2-containing AMPARs into the synaptic membrane by binding to the GluA2 C-terminus (Araki et al. 2010; Huang et al. 2005; Nishimune et al. 1998; Shi et al. 2001). NSF expression from whole cell lysate was unchanged indicating that a key mechanism required for synaptic delivery and activity of GluA2-containing AMPARs was intact in the AcbC following a history of ethanol self-administration. This does not preclude the possibility that the localization of NSF and GluA2 may be disconnected subcellularly, but it still suggests the upregulation of GluA2 and intact NSF protein expression in the AcbC is significant, as we previously observed increased NSF expression in the amygdala of mice drinking ethanol in the home cage but no change in GluA2 (Faccidomo et al. 2021). This suggests that distinct and dynamic glutamatergic mechanisms in the amygdala and AcbC regulate reward-driven behaviors.

These results complement and extend preclinical research showing that home-cage voluntary ethanol drinking alters protein expression of AMPAR GluA2 in the Acb and connected forebrain nuclei. For example, home-cage ethanol drinking increased both GluA1 and GluA2 expression in the synaptic fraction of cells from the dorsal medial striatum of rats at 24-hrs post exposure (Wang et al. 2012) and was associated with an increase in AMPAR GluA1 expression in C57BL/6J mouse Acb at 24-hrs post intake (Ary et al. 2012). Evidence is mixed, however, regarding how long AMPAR subunit adaptations persist in withdrawal and may be brain-region specific. GluA1 expression was not changed in the Acb of rats following extended withdrawal of 40 – 60 days from chronic intermittent ethanol drinking (Marty and Spigelman 2012); however, both GluA1 and GluA2 expression were increased in the prefrontal cortex of male and female C57BL/6J mice following a 30-day history of home-cage ethanol drinking and 28-days of withdrawal (Szumlinski et al. 2023). Together with the results from the present study, these findings present consistent evidence that voluntary ethanol intake is associated with an increase in expression of GluA2-containing AMPARs in reward-related brain regions. Thus, it will be important in future studies to extend our work to evaluate effects of operant ethanol self-administration on AMPAR subunit expression in brain regions where excitatory transmission regulates the reinforcing effects of the drug, including the Acb shell (Valyear et al. 2024), ventral tegmental area (Czachowski et al. 2012) and prefrontal cortex (Faccidomo et al. 2016).

### GluA2-containing AMPAR activity in the AcbC is required for the positive reinforcing effects of ethanol

To address the GluA2 specificity of the AMPAR behavioral regulation of ethanol self-administration via the AcbC, two microinjection studies were conducted. The first used Pep2m which is a cell-permeable inhibitor of GluA2-containing AMPARs that inhibits the interaction between the GluA2 C-terminus and NSF. Disruption of this interaction decreases AMPAR mediated synaptic activity (Nishimune et al. 1998) and surface expression of GluA2-containing AMPARs (Noel et al. 1999). The second used NASPM to inhibit GluA1-containing AMPARs as a test of AMPAR subtype specificity in regulation of ethanol self-administration. Both infusions targeted the AcbC; however, it should be noted that drug solutions may have diffused to adjacent loci.

Intra-AcbC infusion of Pep2m produced a dose-dependent decrease in ethanol - reinforced lever presses, indicating reduced reinforcing effects of ethanol. This led to fewer ethanol reinforcers earned at the highest dose without affecting headpokes in the delivery cup, showing no impact on consummatory behavior. Notably, 10 µg Pep2m reduced response rate during the 20-60 min period of 1-hour sessions without altering the onset of responding. This indicates that Pep2m infusion in the AcbC did not nonspecifically inhibit ethanol operant conditioning or motor ability, as confirmed by an independent open field assessment. Sucrose self-administering mice, used as behavior-matched controls, showed similar responses to the ethanol-trained group under aCSF conditions (∼150 responses/hr). Importantly, however, Pep2m had no effect on sucrose-reinforced responding or spontaneous open field activity, supporting an ethanol-specific effect. These findings highlight the GluA2 and NSF interaction in the AcbC as a novel mechanism of ethanol self-administration.

In this context, it is essential to consider the comparison to the behavior-matched sucrose controls, which is critical to the interpretation that the observed change in GluA2 expression and changes in operant ethanol self-administration were a specific function of ethanol and not related to alternative factors such as general reward processing. The sucrose control also rules out nonspecific intervening behavioral variables such as learning and memory associated with the complex operant task, motor activity, and general consummatory behavior, all of which could be regulated by glutamate activity in the AcbC (e.g., (Grueter et al. 2012; Kelley 2004b)). However, the sucrose control condition does not address potential changes in the pharmacological effects of self-administered ethanol following Pep2m infusion, which could impact ethanol’s discriminative stimulus (Hodge and Cox 1998; Hodge et al. 2001) or rewarding (Pina and Cunningham 2016) properties and reduce self-administration via altered glutamate transmission. Overall, however, these results show that chronic ethanol self-administration produces a reinforcer-specific, AMPAR subunit-selective, increase in GluA2 expression in the AcbC that mediates the reinforcing effects of the drug as definitively measured by a reduction in response rate (Skinner 1938).

GluA2 specificity depends on the brain region. Prior studies have shown that GluA1-containing AMPAR activity in the BLA is required for the reinforcing effects of ethanol (Faccidomo et al. 2021) but it was not known whether GluA1-containing AMPAR activity in the Acb also regulates ethanol’s reinforcing properties. Results of the present study showed that intra-AcbC NASPM had no effect on ethanol self-administration or locomotor activity. This suggests that GluA2-lacking CP-AMPAR activity in the AcbC does not mediate the reinforcing properties of sweetened ethanol as tested under the conditions of the present study. It is worth noting that others have found that NASPM (20 µg/side) infused in the Acb (shell) decreased home-cage ethanol drinking by both female and male C57BL/6J mice with no effect on concurrent water intake (Kwok et al. 2021), suggesting that a higher dosage of NASPM may have shown efficacy in the present study. However, that study used a different ethanol drinking method and higher NASPM dosage suggesting that CP-AMPAR regulation of ethanol operant self-administration and home-cage intake may be Acb-subregion specific and influenced by behavioral factors, such as motivation. Moreover, our prior work showed that the dosage of NASPM (10 µg) used in the present study reduced ethanol self-administration when infused in the amygdala where CP-AMPAR activity was upregulated (Faccidomo et al. 2021). It is also possible that NASPM had no effect on ethanol reinforced responding, compared to Pep2m, due to a lower baseline rate of responding in the NASPM group. This is unlikely, however, since prior studies have shown that a comparable baseline is amenable to treatment-induced decreases or increases in the rate of self-administration (Faccidomo et al. 2020; Faccidomo et al. 2018; Hoffman et al. 2021).Together, these studies underscore the importance of considering neuroanatomical specificity, the nature of the reinforcing or consumed solution, antagonist dosage, and ethanol self-administration method in evaluating AMPAR subtype regulation of ethanol reinforcement.

### Self-administered ethanol increases synaptic activity of CI-AMPARs in AcbC neurons receiving BLA projections

We previously found that ethanol self-administration selectively increased excitatory synaptic activity and membrane insertion of GluA2-lacking CP-AMPARs in BLA neurons projecting to the Acb, and that GluA1-containing AMPARs are required for ethanol’s reinforcing effects (Faccidomo et al. 2021). Here, we extended these findings by examining if operant ethanol self-administration alters synaptic activity of AcbC neurons receiving excitatory BLA projections. Whole-cell patch-clamp recordings from AcbC neurons receiving BLA synaptic input showed that chronic ethanol self-administration (X̅ = 0.79 g/kg/day) increased sEPSC decay time and area under the curve, indicating a shift to increased synaptic representation of GluA2-containing AMPARs (Chater and Goda 2014; Pampaloni and Plested 2022), consistent with upregulated GluA2 protein expression in the AcbC and microinjection results.

To determine the impact of ethanol self-administration on synaptic activity, we recorded glutamatergic transmission with whole-cell patch clamp recordings in the BLA→AcbC circuit and found that optical stimulation of ChR2-positive BLA terminals in the AcbC exhibited a strong trend (P = 00518) in the oEPSC AMPA/NMDA ratio in ethanol self-administering mice as compared to sucrose controls suggesting an ethanol-induced increase in synaptic strength or plasticity, or a potential increase in the number or function of AMPARs, which is also consistent with our observation of upregulated GluA2 protein expression. Interestingly, the AMPA/NMDA ratio was not altered following local electrical stimulation in the AcbC suggesting an ethanol-specific effect on BLA terminals. While the current study did not differentiate AcbC cell types, qualitatively examining the data in the cells from ethanol self-administering animals, the AMPA/NMDA ratios clustered into 3 distinct groups, suggesting that there may be differences in BLA plasticity onto distinct cell types in the AcbC. We plan to explore this possibility in future studies. In addition, we observed a significant decrease in the paired-pulse ratio (PPR) in ethanol self-administering mice as compared to sucrose controls following optical stimulation of ChR2-positive BLA terminals in the AcbC. While one interpretation of paired-pulse depression (PPD) of oEPSCs is indicative of an increase in the probability of glutamate release, another interpretation of PPD is a change in synaptic subunit composition to synapses with increased GluA2 expression (Liu and Savtchouk 2012). Although the specific mechanism of PPD was not pursued in this study, it is likely influenced by enhanced synaptic activity GluA1-containing CP-AMPARs in the BLA projection neurons as we discovered previously (Faccidomo et al. 2021), and may be modified by a variety of related cellular processes including increased CaMKII expression or activity in the BLA, or upregulation of AMPAR accessory proteins such as TARP ɣ-8 in the postsynaptic membrane, both of which enhance AMPAR activity (Hayashi et al. 2000; Tomita et al. 2003) and regulate ethanol self-administration (Agoglia et al. 2015; Cannady et al. 2016; Faccidomo et al. 2016; Hoffman et al. 2021; Hoffman et al. 2023; Salling et al. 2016).

Finally, bath application of the CP-AMPAR inhibitor NASPM confirmed the specificity of GluA2-containing AMPAR activity on synaptic function. NASPM had no effect on oEPSC amplitude in either ethanol or sucrose self-administering mice, suggesting the absence of a significant population of CP-AMPARs in the BLA → AcbC synapses following the operant behavioral protocol used in this study. This contrasts with our previous discovery in BLA neurons projecting to the Acb, where NASPM selectively decreased eEPSC amplitude in ethanol self-administering mice but not in sucrose controls, indicating increased membrane insertion of CP-AMPARs in the BLA (Faccidomo et al. 2021). Together, these studies support the conclusion that ethanol self-administration produces differential, brain-region-dependent effects on AMPAR subunit composition and synaptic function, with upregulation of GluA2-containing CI-AMPAR activity in the AcbC and GluA1-containing AMPAR activity in the BLA. These data also suggest increased plasticity in both pre- and postsynaptic compartments in the BLA → AcbC circuit following ethanol self-administration.

### Overall Conclusions

The implications of these experiments are significant for understanding how neural plasticity within the BLA → AcbC excitatory circuit regulates the positive reinforcing properties of ethanol. Numerous reports indicate that GluA2-lacking CP-AMPARs, which can be GluA1 homomers, are rapidly inserted in synapses undergoing specific forms of synaptic plasticity, such as long-term potentiation, and following tests of behavioral plasticity, including fear conditioning and exposure to drugs of abuse (reviewed by (Diering and Huganir 2018)). However, the synaptic expression of CP-AMPARs is transient and followed by replacement with GluA2-containing CI-AMPARs (Plant et al. 2006), stabilizing long-term changes in synaptic efficacy (Shi et al. 2001). Our prior results showing upregulated CP-AMPAR activity in BLA neurons projecting to the Acb following chronic ethanol self-administration (Faccidomo et al. 2021) suggest ongoing plasticity in BLA projection neurons with repeated ethanol exposure. Alternatively, AcbC neurons receiving BLA projections exhibit increased GluA2-containing CI-AMPAR expression and activity following chronic ethanol self-administration, consistent with long-term stabilization of ethanol’s impact on neural function in this key reward system nucleus (Plant et al. 2006). These observations raise the hypothesis that GluA1-containing CP-AMPAR activity in the BLA may drive ongoing changes in ethanol’s reinforcing effects, whereas GluA2-containing CI-AMPAR activity in the AcbC may underlie the long-term maintenance, or memory, of this maladaptive stimulus property of the drug. It is a limitation of this study, however, that we used only male mice, which narrows the generality of the findings. This approach maintains consistency with our prior research in the amygdala (Faccidomo et al, 2021), allowing direct comparisons. Additionally, existing literature suggests that female mice self-administer ethanol at higher levels than male mice (Sneddon et al, 2020), which introduces differential ethanol intake as confounding variable when evaluating the effects of voluntary self-administration on brain function. Nonetheless, the importance of evaluating sex as a biological variable (SABV) cannot be overstated, as physiological and neurobiological responses often differ between sexes, potentially impacting both addiction pathways and behavioral outcomes (Becker and Koob, 2016). Future studies are essential to address potential sex differences in AMPAR regulation of the reinforcing effects of ethanol, ultimately ensuring a more comprehensive understanding of excitatory neural circuit function in alcohol-related behaviors across both male and female populations.

In conclusion, harmful alcohol use is a significant public health concern worldwide (WorldHealthOrganization 2018), and understanding the neural mechanisms underlying its positive reinforcing effects is crucial for developing effective treatments (Stolerman 1992). Here we show evidence that ethanol targets AMPAR GluA2 expression and activity in the Acb, a key nucleus in the brain’s reward pathway, regulating the drug’s reinforcing effects. Based on these results and observations that long-term memory consolidation requires GluA2-containing AMPARs (Pereyra and Medina 2021), we propose that ethanol’s ability to drive repetitive self-administration in AUD is mediated through hijacking this fundamental process in the Acb. Furthermore, pharmacological tools like Pep2m, which mediate AMPAR GluA2 endocytosis and downregulation, have potential to promote forgetting of specific long-term memories (Hardt et al. 2014; Migues et al. 2016), such as the reinforcing effects of alcohol. Future studies should test this hypothesis using models of cue-induced reinstatement and other techniques that specifically test the regulation of alcohol memories via AMPAR activity.

